# Cell Cycle Phases, Spindle Dynamics and Kinesin-5 Motor Localization Characterized by Deep Learning, Dual Segmentation and Decision-Tree Pipeline

**DOI:** 10.64898/2026.08.24.746832

**Authors:** Omer Bushusha, Karin Zarnitsky, Neta Yanir, Mayan Sadan, Daniel Sevilla Sánchez, Larisa Gheber

**Affiliations:** Department of Chemistry and Technology, Ben-Gurion University of the Negev, Beer-Sheva, Israel; Ilse Katz Institute for Nanoscale Science and Technology, Ben-Gurion University of the Negev, Beer-Sheva, Israel

## Abstract

Three-dimensional live-cell fluorescence imaging of yeast cells is crucial for studying cell-cycle mechanics and regulation. However, extracting multi-channel phenotypes within dense cell clusters remains an image-processing bottleneck. Standard deep-learning models segment cells but fail to track mother-bud boundaries, mitotic spindle shapes and spindle-localizing proteins. Investigators rely on labour-intensive manual coordinate plotting, introducing observer bias and often exclude clustered cell data due to visual complexity. Here, we present an open-source Fiji pipeline for automated yeast cell image processing and deterministic classification of cell-cycle, spindle and protein dynamics. The workflow utilizes a dual-segmentation architecture via custom Cellpose models to capture the mother-bud cell boundaries. Extracted masks are integrated with multi-channel fluorescence data using a Difference-of-Gaussians framework to resolve SPB coordinates and localized protein kinetics, which a rule-based decision-tree maps to precise mitotic phenotypes. Validation demonstrates a 50-fold acceleration with ∼6% deviation from manual analysis.

**Availability:** Zenodo at https://doi.org/10.5281/zenodo.22083016.

## Introduction

In eukaryotic cells, the orchestration of mitosis requires the precise spatiotemporal regulation of the mitotic spindle, an anisotropic macromolecular machine composed of microtubules (MTs) and specialized spindle-binding proteins (Reviewed in (Goulet and Moores, 2013; Howard and Hyman, 2003; Kapoor, 2017; Liakopoulos, 2021; Mann and Wadsworth, 2019; Nazockdast and Redemann, 2020; Scholey, et al., 2003; Wadsworth, 2021)). Spindle assembly, positioning, and elongation are driven by a delicate balance of forces generated by kinesin and dynein motor proteins (Heald, 2000; Pandey, et al., 2021; Sharp, et al., 2000; Singh, et al., 2018) and by MT dynamics(Dumont and Mitchison, 2009; Mitchison and Kirschner, 1984). The budding yeast *Saccharomyces cerevisiae* serves as an excellent model for studying cell cycle progression and mitosis. In these cells, structural dynamics of the spindle occur concurrently with morphogenetic changes, such as the emergence and growth of the daughter bud and the positioning of the nucleus at the mother-bud region. Moreover, for correct mitotic progression, several regulatory and functional proteins change their localization along the mitotic spindle in a spatiotemporally regulated, dynamic manner. These include non-motor MT crosslinking proteins (Alfieri, et al., 2021; Fu, et al., 2009; Gaska, et al., 2020; Schuyler, et al., 2003) chromosomal passenger complexes (Carmena, et al., 2012; Hadders and Lens, 2022; Ibarlucea-Benitez, et al., 2018) and kinesin motor proteins(Goldstein-Levitin, et al., 2021; Goldstein, et al., 2017; Mann and Wadsworth, 2019; Singh, et al., 2024). Thus, tracking these cellular and subcellular morphological changes serves as a vital tool for understanding cell-cycle progression, mitotic regulation, and force-balance kinetics at the single-cell level.

Consequently, high-content, three-dimensional (3D) fluorescence live-cell imaging has become an indispensable tool for characterizing how specific genetic mutations or biochemical perturbations disrupt these mechanical processes (Stephens and Allan, 2003). However, a severe bottleneck persists at the stage of image processing and phenotypic extraction. While deep-learning frameworks such as U-Net(Falk, et al., 2019), StarDist(Schmidt, et al., 2018), and Cellpose (Pachitariu and Stringer, 2022; Stringer, et al., 2021) have significantly improved individual cell segmentation boundaries, mapping complex, multi-channel structural phenotypes remains highly problematic. In budding yeast, this challenge is doubled: a meaningful phenotypic readout requires the simultaneous segmentation of individual cell boundaries, the identification of the specific mother-bud cell pair, and the localization of internal structures like spindle pole bodies (SPBs) and localized, spindle-bound proteins. Modern platforms often require investigators to construct post-segmentation scripts or perform labor-intensive manual pedigree tracking, which restricts workflow scalability, introduces possible observer bias, and lacks integrated validation interfaces capable of combining spatial data across brightfield and fluorescent channels(Padovani, et al., 2022).

This limitation is particularly pronounced when tracking sub-micron structures or analyzing densely packed cell clusters. During early spindle assembly in *S. cerevisiae* cells, when duplicated SPBs are separated by short distances (L < 1.8 µm), overlapping fluorescence emission profiles can lead to subjective, inconsistent manual coordinate selections, potentially leading to overestimation of these short distances, artificially overestimating the scores of longer spindles in the anaphase spindle elongation phase (Goldstein, et al., 2017; Higuchi and Uhlmann, 2005; Movshovich, et al., 2008; Straight, et al., 1998). Consequently, this leads to the underestimation of cells with short-bipolar spindles, as well as spindle elongation delays characteristic of cells carrying mutations in proteins regulating the metaphase-to-anaphase transition(Cohen-Fix and Koshland, 1997; Severin, et al., 2001; Stemmann, et al., 2001). Furthermore, human annotators routinely discard densely packed cell regions due to the visual complexity of manual tracing, thereby discarding biological data and introducing systematic bias toward isolated single cells.

To bridge the gap between automated pixel segmentation and biological phenotype classification, we present the Spindle-Phenotype Characterization (SPC) tool, a Fiji-based (Schindelin, et al., 2012) software pipeline engineered for high-throughput, multi-channel *S. cerevisiae* cell analysis. Our approach integrates a specialized dual-segmentation strategy using Cellpose (Pachitariu and Stringer, 2022; Stringer, et al., 2021) to concurrently capture independent cell masks and mother-bud cell boundaries. By extracting deterministic physical measurements such as Difference-of-Gaussians (DoG)-derived SPB coordinates, automated tracking of variation in spindle-bound protein localization, in this case, kinesin-5 mitotic spindle motor protein Cin8 (Gerson-Gurwitz, et al., 2011; Goldstein-Levitin, et al., 2021; Hildebrandt and Hoyt, 2000; Saunders and Hoyt, 1992), and exact bud-to-mother diameter ratios, this tool passes each cell through a rigid, traceable decision tree. The pipeline automatically categorizes mitotic phenotypes, including cells with monopolar, short bipolar, and early/late anaphase spindles, as well as the telophase stage. By offering both an automated execution and an active, online human validation interface (human-in-the-loop), this tool eliminates human spatial bias, analyzes clustered cell-populations otherwise omitted from manual workflows, and accelerates analysis throughput by approximately 50-fold, providing an accurate and reproducible platform for large-scale phenotypic profiling of the *S. cerevisiae* mitotic machinery.

## Experimental Methods

Live cell imaging was performed as previously described, using *S. cerevisiae* strains expressing tdTomato-labeled spindle pole body (SPB)-binding protein Spc42(Pandey, et al., 2021). In addition, cells expressed one 3GFP-labeled kinesin-5 variant, Cin8 or Kip1. Cin8-3GFP expressing strain is: (LGY3989: MATa, ura3-52, leu2-3,112, his3-Δ200, lys2-801, cin8::LEU2, spc42::KanMX-SPC42-TdTomato (pVF68: CEN, URA3, CIN8-3GFP)). Cells were grown in selective medium, till mid-log phase, followed by placement of a sample of cells on a low-fluorescence agarose gel on a slide. Images were acquired at room temperature using a Nikon Ti2 inverted fluorescence microscope controlled by NIS-Elements AR (v6.10.01). The system included an Andor Zyla sCMOS camera and a Plan Apo λD 100×/1.45 NA oil DIC objective. Illumination was provided by a CoolLED pE-800, and fluorescence was recorded with 515/30 (GFP) and 595/31 (tdTomato/TRITC) emission filters. Z-stacks of 30 planes were acquired with ∼0.30 μm spacing.

### SPC Tool Development

#### Data Preparation and Processing

The developed tool processes three-channel 3D image stacks: brightfield (cell shape), green fluorescence (spindle-bound GFP-labeled kinesin-5 motors) and red fluorescence (tdTomato-labeled SPBs); and projects them into 2D for signal strength and contrast enhancement, and robust segmentation and feature extraction. The use of 2D projections is appropriate because cells in the colonies are arranged laterally rather than axially and are therefore rarely stacked on top of each other (Fig. 1A-C). Following 2D projection, all channel projections are normalized, stretching their dynamic range to the full 16-bit dynamic range (0-65535) before Cellpose training and inference to standardize intra-channel intensity ratios, improve visibility, and minimize variability between datasets. Fluorescence channels are projected using maximum-intensity projection. For the brightfield channel, the pipeline computes both maximum- and minimum-intensity projections and subtracts the minimum from the maximum (Fig. 1D-F). This differential projection markedly improves local contrast, suppresses background, and yields superior performance with the default Cellpose models.

**Figure 1.**
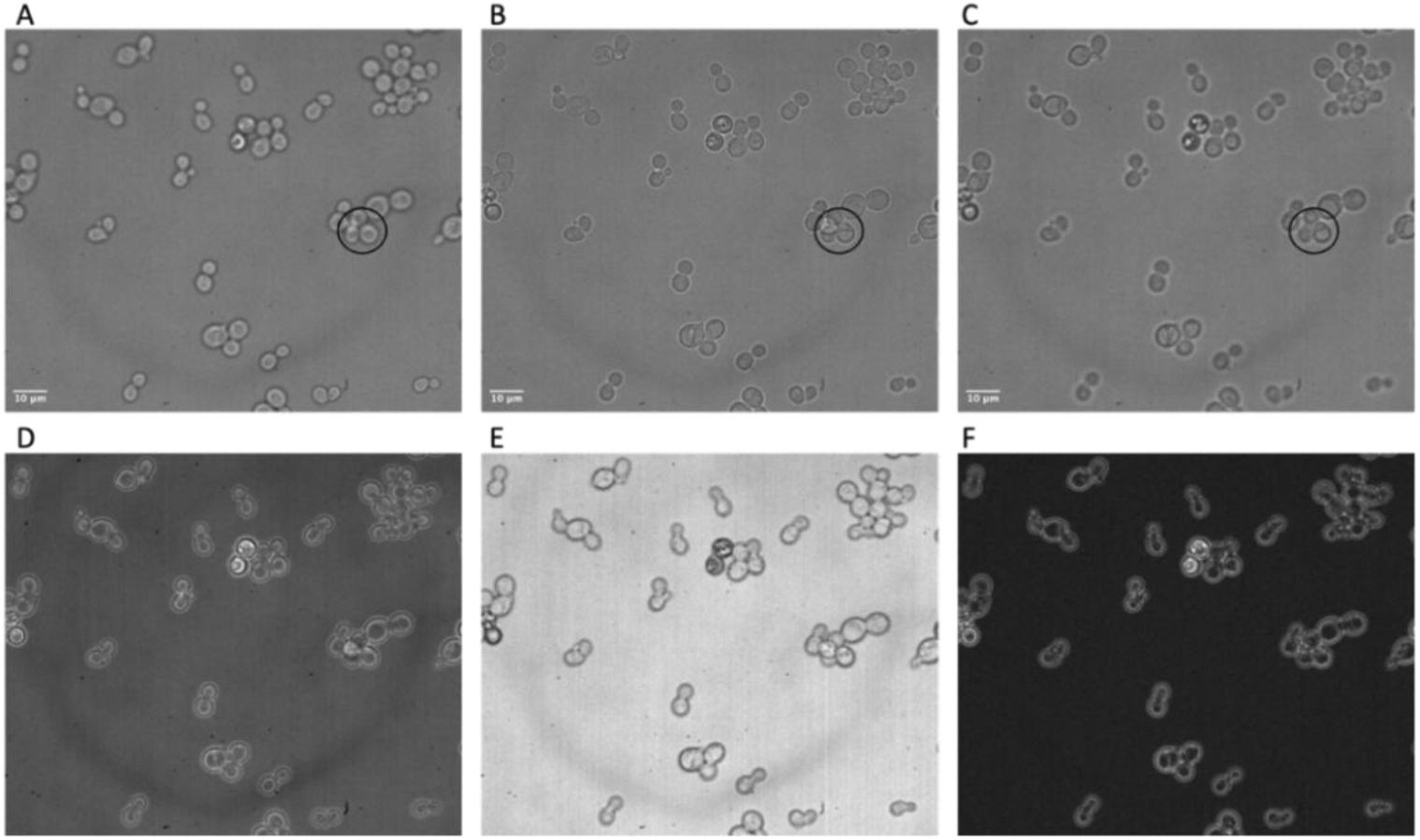
Data preparation for the SPC tool. **A-C** bright-field images of *S. cerevisiae* cells captured at three Z-planes (1, 10, and 15, respectively) out of total 30 planes. The circled region is an example of rare axial overlap. **D and E** are the maximum and minimum intensity projections of the same field, respectively. **F** field resulting from subtracting the minimum intensity projection from the maximum intensity projection. Bar: 10 µm.

#### Segmentation

Segmentation of cells is performed on the normalized three-channel 2D projections. To enhance prediction accuracy, we employed a transfer-learning strategy, starting from the default Cellpose (Pachitariu and Stringer, 2022; Stringer, et al., 2021) models and iteratively fine-tuning them with curated corrections and additional annotations provided by lab members experienced in *S. cerevisiae* cell division. Two specialized models were trained for distinct but complementary tasks (Fig. 2):

**Figure 2.**
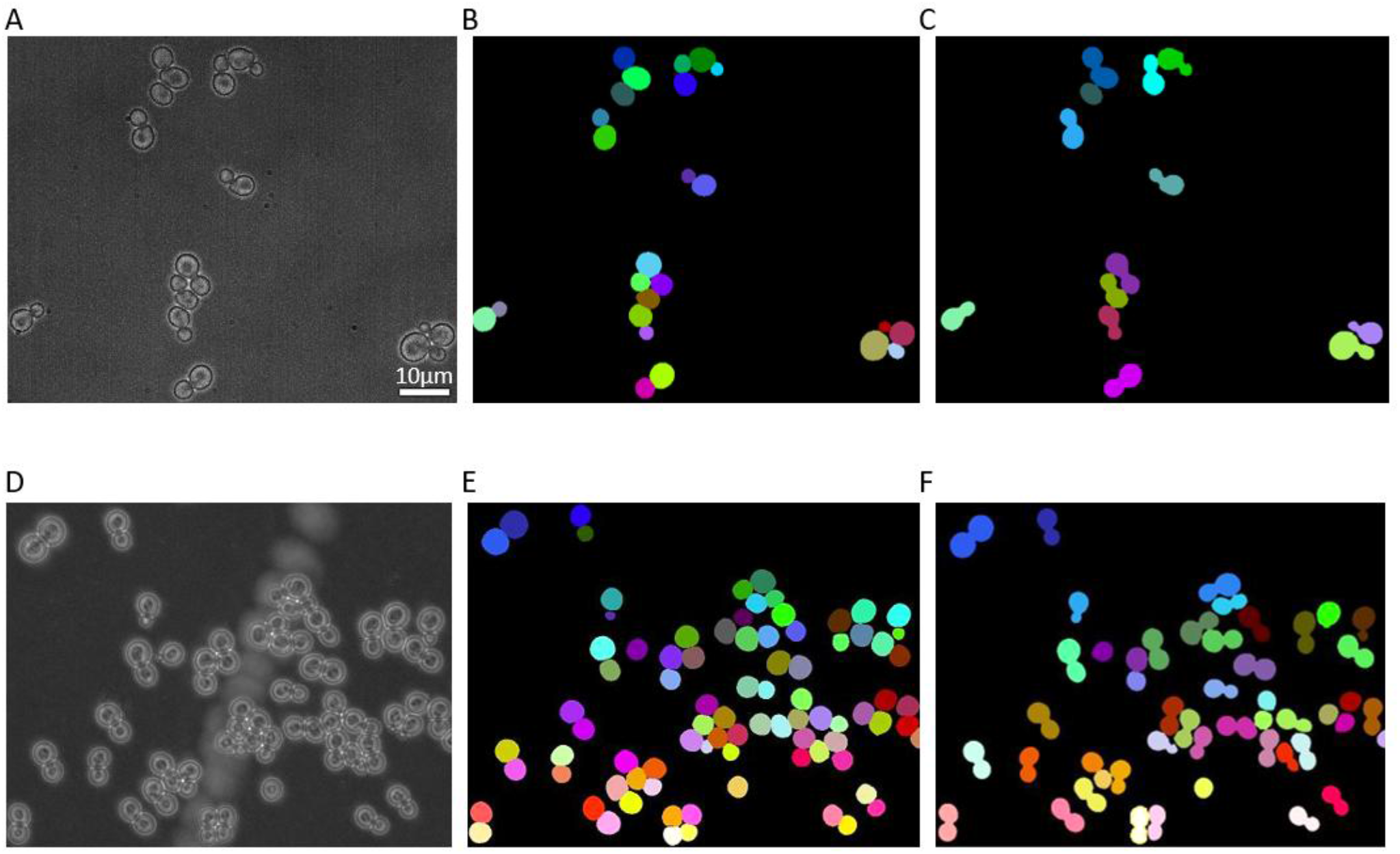
Models used by the SPC tool. **A and D**, bright-field images of *S. cerevisiae* cells from a live-cell imaging assay expressing GFP-labeled Cin8, respectively. The images are the result of subtracting the minimum-from the maximum-intensity projection. **B and E**, the ‘allObjectsImg’ model identifies all cells as masks. **C and F**, the ‘cellLabelsImg’ model identifies mother-bud pairs as masks. Bar: 10µm.

1. A compound model for detecting mother-bud pairs, enabling recognition of connected mother-bud systems.
2. A single-cell model for identifying individual cells, mothers and buds, as independent objects.

Combining the outputs of both models enhanced coverage across a range of morphological configurations, from isolated cells and mother-bud pairs to large, complex clusters containing multiple touching individual cells and mother-bud pairs. The two models exploit complementary structural information from the bright-field and fluorescence channels, including cell morphology and boundaries, spindles and spindle poles. This information is particularly useful for resolving cells within crowded clusters, where the spindle provides a directional cue that can help identify which adjacent cells form a mother-bud pair. The identification of mother-bud pairs in budding yeast has also recently been addressed (Zhao, et al., 2026).

#### Detection of SPBs and motor protein spindle localization

The trained Cellpose models form the basis for accurate cell segmentation and subsequent phenotype classification. Users supply the three-channel images; the tool then performs the 2D projections and invokes the two Cellpose models through the PTBIOP plugin in Fiji^1^. These sequential calls generate two label images:

1. Mother-bud objects.
2. Independent single-cell objects.

The mother-bud labels are imported into the Region Of Interest (ROI) Manager in Fiji via the PTBIOP plugin. The tool iterates over these ROIs, processing each mother-bud object to measure geometric features and to generate per-cell thumbnails (Fig. 3, bottom panel of each cell morphology and spindle phenotype). All resulting data and images are automatically written to the specified output directory.

**Figure 3.**
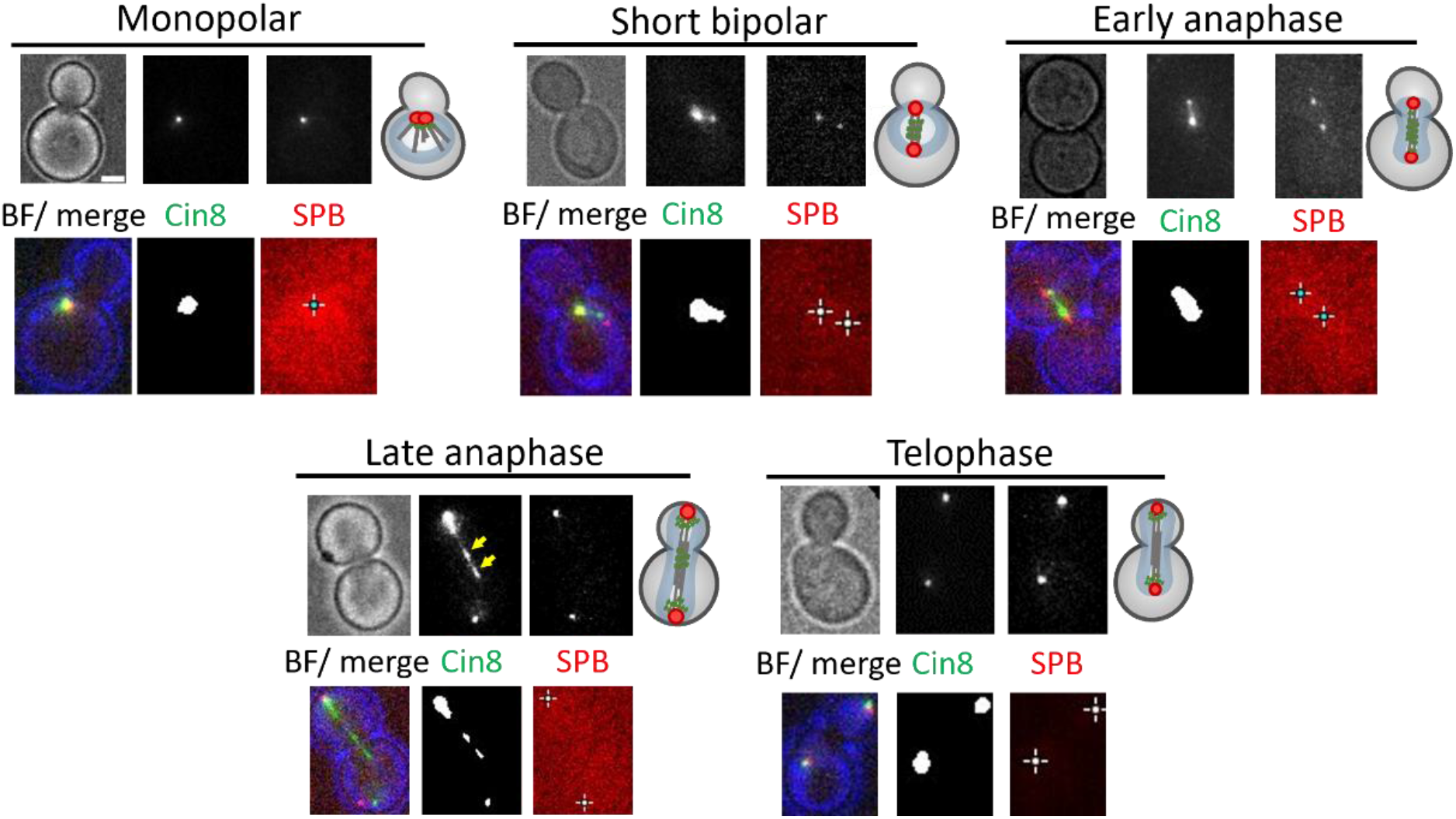
SPBs and Cin8 (MSP) detection. Localization of Cin8-3GFP variants in *S. cerevisiae* cells expressing tdTomato-labeled SPB component Spc42. Bright field (BF), Cin8 and SPBs images are shown. Schematic representation of budded cells with spindles of different morphologies is shown on the left of each panel. **Top of every phenotype** – images acquired by the microscope. These cells were sometimes rotated for representation purposes. **Bottom of every phenotype** - output images from the SPC tool for the same cells, representing the location of the SPBs and Cin8 GFP signal detected by the tool. These images were not rotated. Spindle morphologies are categorized as follows: monopolar spindles – 1 SPB (the red signal of the two SPBs are indistinguishable); short bipolar (pre-anaphase) spindles – L < 1.8 µm; intermediate bipolar (early anaphase) spindles – 1.8 µm < L < 4 µm; long bipolar (late anaphase) spindles – 4 µm < L; telophase spindles – 3 µm < L with GFP signal concentrated near the SPBs. Yellow arrows point to GFP puncta; Bar – 2 µm.

SPB detection is implemented as a robust, multi-step procedure:

1. Extract the red-channel ROI.
2. Apply a difference-of-Gaussians (DoG) sweep across multiple σ values to identify candidate blobs.
3. For each DoG candidate, locate local maxima.
4. Around each maximum, compute a robust Z-score within a disk-ring system centered on the maximum.
5. Select the two peaks with the highest Z-scores exceeding a conservative threshold.

The disk-ring Z-score quantifies how many robust standard deviations (computed using the median absolute deviation (MAD), to discard outliers) the inner-disk intensity exceeds that of the surrounding ring, with the ring median serving as the reference and the MAD as a robust spread estimate. After empirical threshodkinesin and dynein motor proteins l tuning, this method achieves high SPB-detection accuracy even under low signal-to-noise conditions.

Detection of kinesin-5 motor protein localization is performed by extracting the green-channel ROI and assessing the presence of GFP signal. Background is subtracted, followed by a Gaussian smoothing (σ = 1). From the filtered ROI, the coefficient of variation (CV) and contrast ratio (CR) are computed. Conservative thresholds on CV and CR exclude structureless regions, while refined thresholds distinguish weak diffuse signal from true motor-localization structures. For simplicity, we refer to the motor protein GFP localization as “mitotic spindle-binding protein” or MSP. When this MSP GFP signal is detected, the region is segmented using Yen thresholding (Yen, et al., 1995), an automatic histogram-based method for choosing the intensity cutoff, with objects smaller than 4 pixels (~0.5 µm) discarded.

To determine whether the MSP extends from and between the detected SPBs, which is necessary for detection of Early and Late Anaphase, the tool combines SPB and GFP MSP data while avoiding misalignment artifacts. The SPBs are temporarily replaced by two 1.5 μm-diameter disks to mask their immediate vicinity; a bounding box enclosing these disks is defined. If the MSP signal remains within this box without overlapping the SPB disks, the MSP is considered consistent, as in Early Anaphase morphology, for example.

We have previously reported that in some instances, Cin8 and Kip1 motor proteins exhibit puncta-like localization to the mitotic spindles, which is related to the functions of these motors (Fridman, et al., 2013; Goldstein, et al., 2017). Thus, the tool was expanded to detect motor protein spindle localization in puncta. Puncta detection reuses the segmented MSP as a mandatory mask. The green-channel ROI is processed with a DoG filter; the result is subjected to thresholding using the Yen method and filtered to exclude objects smaller than 5 pixels (0.64 µm). This is combined with a mask containing the five highest local maxima in the DoG image to improve precision. Puncta located within 0.75 μm near the SPBs are excluded from analysis.

#### Decision-tree-based phenotype classification

Cell-cycle and mitotic phenotypes for each cell are classified using a rule-based decision tree that relies exclusively on measured physical parameters. The main cues are presence and size of the bud, the number of SPBs, pole-to-pole distance, presence or absence of a GFP MSP signal between SPBs, and GFP puncta count. The phenotype decision rules are defined as follows:

- 1 SPB → Monopolar spindle
- 2 SPBs →
  - 0 - 1.8 μm pole-to-pole distance → Short Bipolar spindle
  - 1.8 - 4 μm pole-to-pole distance →
- GFP MSP signal between the SPBs is detected → Early Anaphase
- GFP MSP signal between the SPBs is not detected → Telophase
  - 4 μm pole-to-pole distance →
- GFP MSP signal between the SPBs is detected→ Late Anaphase
- GFP MSP signal between the SPBs is not detected → Telophase

For each cell, the tool reports the phenotype label, puncta count, GFP MSP signal presence or absence in the green-channel, number of SPBs, pole-to-pole distance, geodesic length of the largest MSP component, and bud type (none, small, or large).

For classification of the bud type, we combine information from both Cellpose outputs. The ROI corresponding to a mother–bud detection is compared with the label image containing the independently segmented cells to identify which individual cell labels, if any, overlap the corresponding mother-bud ROI. Using MorpholibJ (Legland, et al., 2016), area and circularity are measured for each component. If two distinct regions cannot be resolved by this method, then the ROI is re-examined for shape indentation, which typically indicates a bud; a binary watershed then separates the ROI into two plausible components, and their areas are measured. Bud size is assigned by comparing the equivalent diameters of both regions:

- Mother diameter > 2× Bud diameter → Small bud
- Mother diameter < 2× Bud diameter → Large bud

All classification outcomes, including intensity and geometric measurements, are compiled into a summary table. The tool also generates thumbnails of each cell and channel, as well as full field-of-view screenshots for visual reference and validation (Fig. 3).

### Output generation

The phenotype-assignment pipeline produces three complementary types of output:

1. Annotated thumbnail images for each detected cell (Fig. 3, bottom panel of each phenotype), facilitating rapid visual inspection and validation.
2. Comprehensive measurement tables containing both raw and derived features, including geometric, and classification data.
3. Metadata and processing logs documenting all parameters and analysis steps to ensure full reproducibility.

All outputs are organized for direct integration into downstream statistical analysis and visualization workflows, enabling seamless coupling with external platforms such as Fiji, R, or Python-based pipelines.

### User correction and validation

During iterations over the detected cells, the tool highlights each ROI and prompts the user for confirmation, enabling rapid triage based on the presence of SPBs, GFP MSP signal, and buds. After all detected cells have been reviewed, users can correct missed pairings by clicking on the object (cell) label image to designate adjacent objects as mother-bud pairs. These manually defined pairs are analyzed using the same pipeline as the automatically detected ones, ensuring consistency in feature extraction and classification. To further improve coverage, a drawing mode allows users to manually outline missing cells directly on the three-channel 2D projection. These user-defined ROIs are then subjected to the same measurement and classification workflow described above. For datasets with high signal quality and reliable automatic detections, the tool also provides a fully automatic mode, which skips interactive prompts and correction steps while producing identical per-cell and per-field-of-view outputs.

To evaluate the accuracy of the SPC tool in classifying cells into phenotypes, we analyzed three colonies of *S. cerevisiae* expressing WT Cin8 (Fig. 4). This dataset was selected since it had been analyzed manually in our previous study (Pandey, et al., 2021). The distributions of cell cycle and spindle phenotypes obtained with and without user correction were compared to assess consistency. Two primary sources of classification errors were identified: SPB detection and GFP motor detection (Fig. 4A). Errors in SPB detection typically resulted in cells being misclassified as monopolar instead of short-bipolar (Fig. 4B). In contrast, errors in detecting spindle-associated motors led to cells being categorized as early or late anaphase rather than telophase. Both error types stem from the chosen detection threshold; however, lowering this threshold would increase false-positive outputs, which we aimed to avoid. Motor misdetections occurring in about 11% of cells were easily corrected within the tool’s interface and required minimal additional time. SPB misdetections occurred in approximately 6% of cells and required manual correction by measuring pole-to-pole distance. Overall, 6% of phenotype assigning errors are minimal, resulting in similar phenotype distributions between corrected and non-corrected tool application (Fig. 4B).

**Figure 4.**
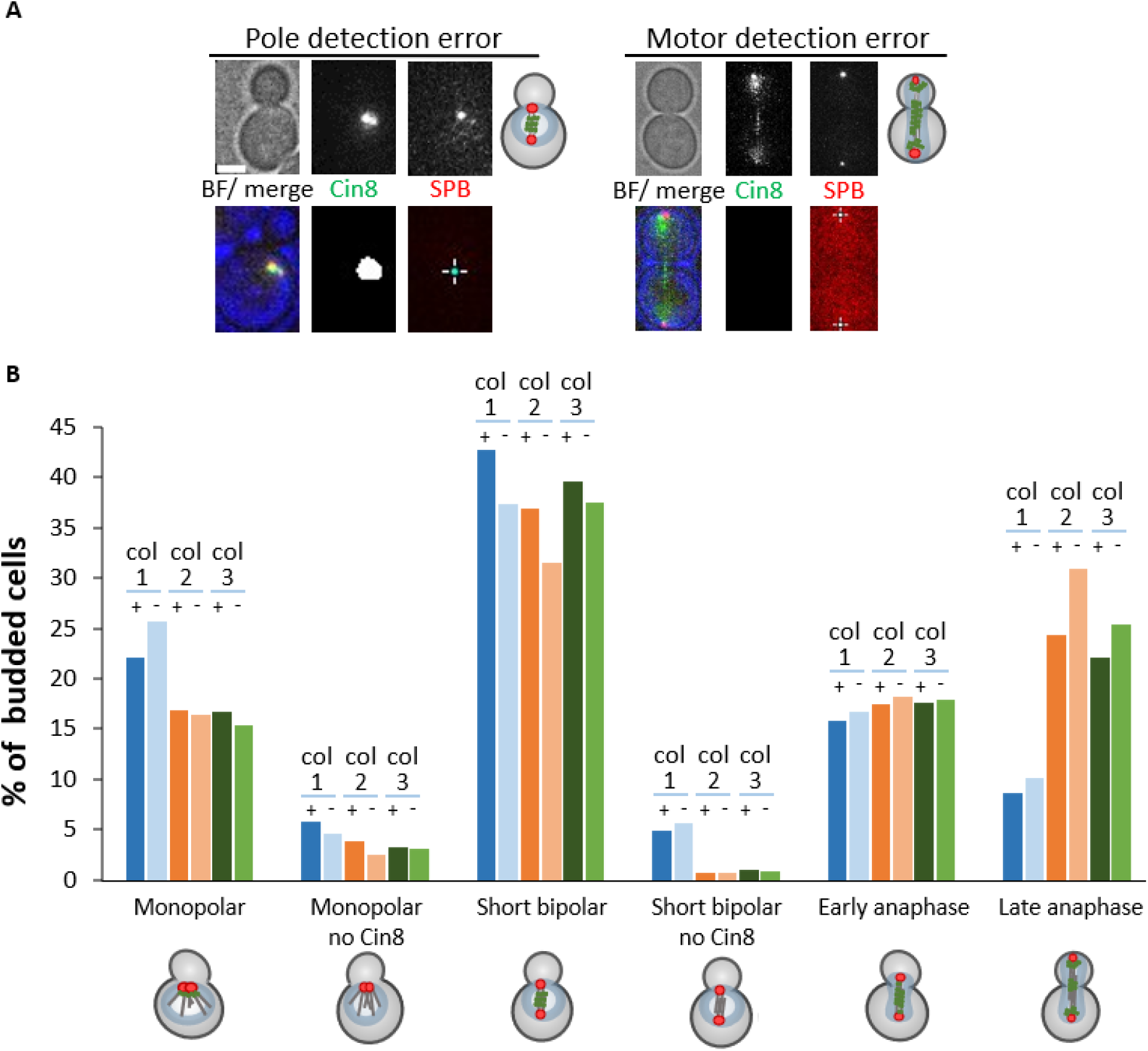
Tool-assigned cell cycle and spindle phenotypes, with and without user correction. **A** Two main types of detection errors by the SPC tool, misdetection of SPB tdTomato signal (left), or misdetection of motor GFP signal (right). The top images are acquired by the fluorescence microscope, the bottom images are outputs from the SPC tool for the same cells, representing the location of the SPBs and Cin8 signal detected by the tool. **B** Cell cycle and spindle phenotype distribution for three independent yeast colonies expressing WT Cin8, with and without user corrections. Each bar represents a specific phenotype of a single colony, shown both in the presence (+) and absence (-) user correction. For each colony, 250 budded cells were categorized. Schematic representations of budded cells with spindles of different lengths and morphologies are shown.

### Comparison between the SPC tool-mediated and manual phenotype classification

The automated analysis pipeline was compared to manual and user-corrected analyses using datasets from three *S. cerevisiae* colonies, published previously by us (Pandey, et al., 2021) (Fig. 5). Processing with the SPC tool was approximately 50-fold faster than manual analysis, reducing the analysis time from weeks and even months to just 1-2 days per colony. Comparison of phenotype classification results revealed no significant differences between the corrected and non-corrected analyses, and only one instance of significant deviation when compared to manual analysis (Fig. 5). We checked whether the primary reason for this discrepancy may be the improved ability of the SPC tool to accurately segment and analyze cells within clusters, a task that was often not achievable during manual processing. Notably, approximately 50% of the cells analyzed using the SPC tool were part of such clusters. To check if the analysis including cells in clusters harms classification accuracy, we compared the data that was acquired by the tool with and without cells that are classified as in clusters (Fig. 6) and saw no significant differences between them.

**Figure 5.**
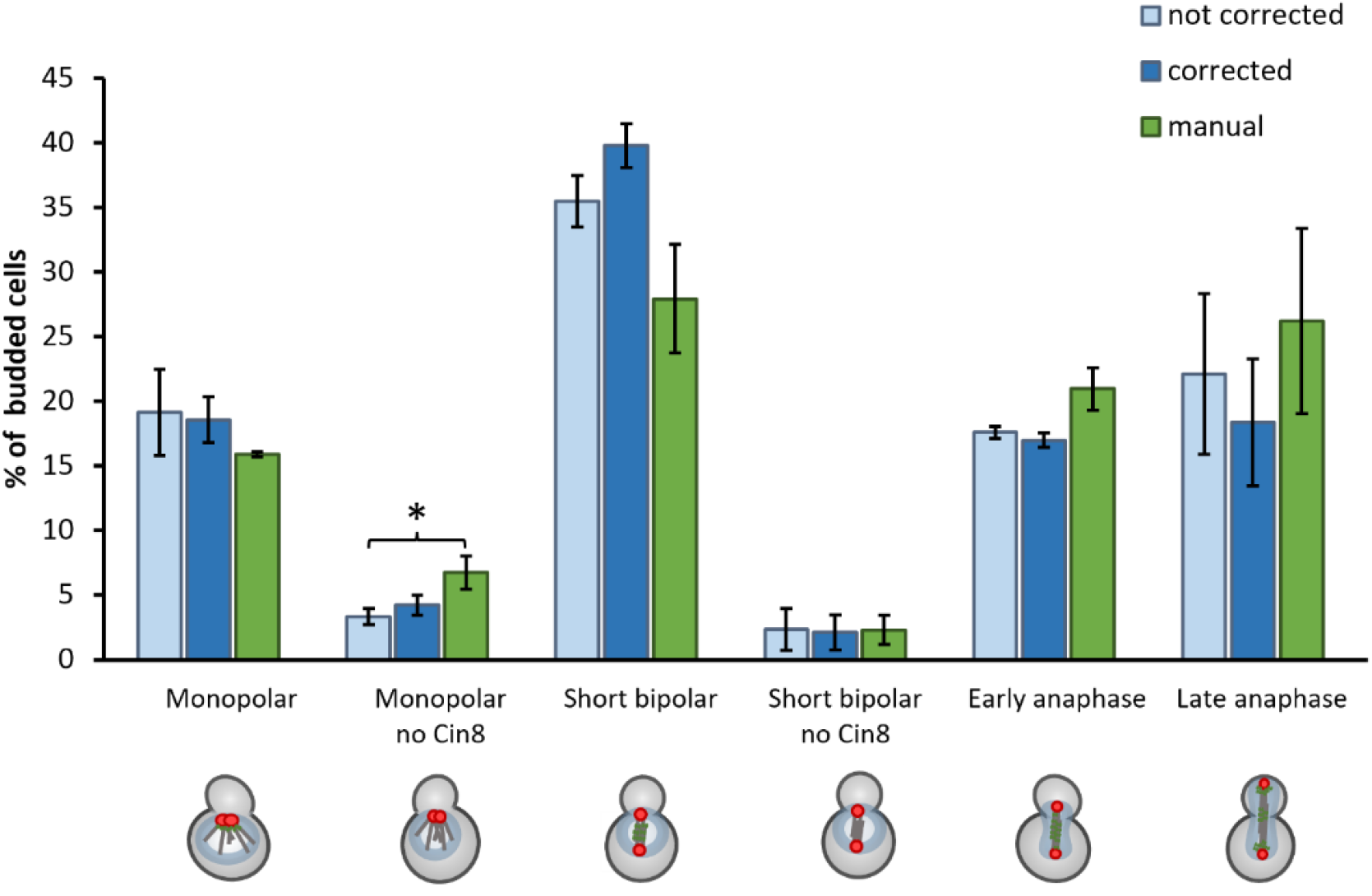
Distribution of Cin8 localization phenotypes. Comparison between three analyses: SPC-mediated, not corrected (light blue), SPC-mediated, user corrected (dark blue) and manual (green). Columns and bars represent averages (± SEM) of 3 independent experiments. \**P* < 0.05; calculated for each spindle morphology, by Student’s t-test. Only significant comparisons (*P < 0*.*05*) are marked with an asterisk.

**Figure 6.**
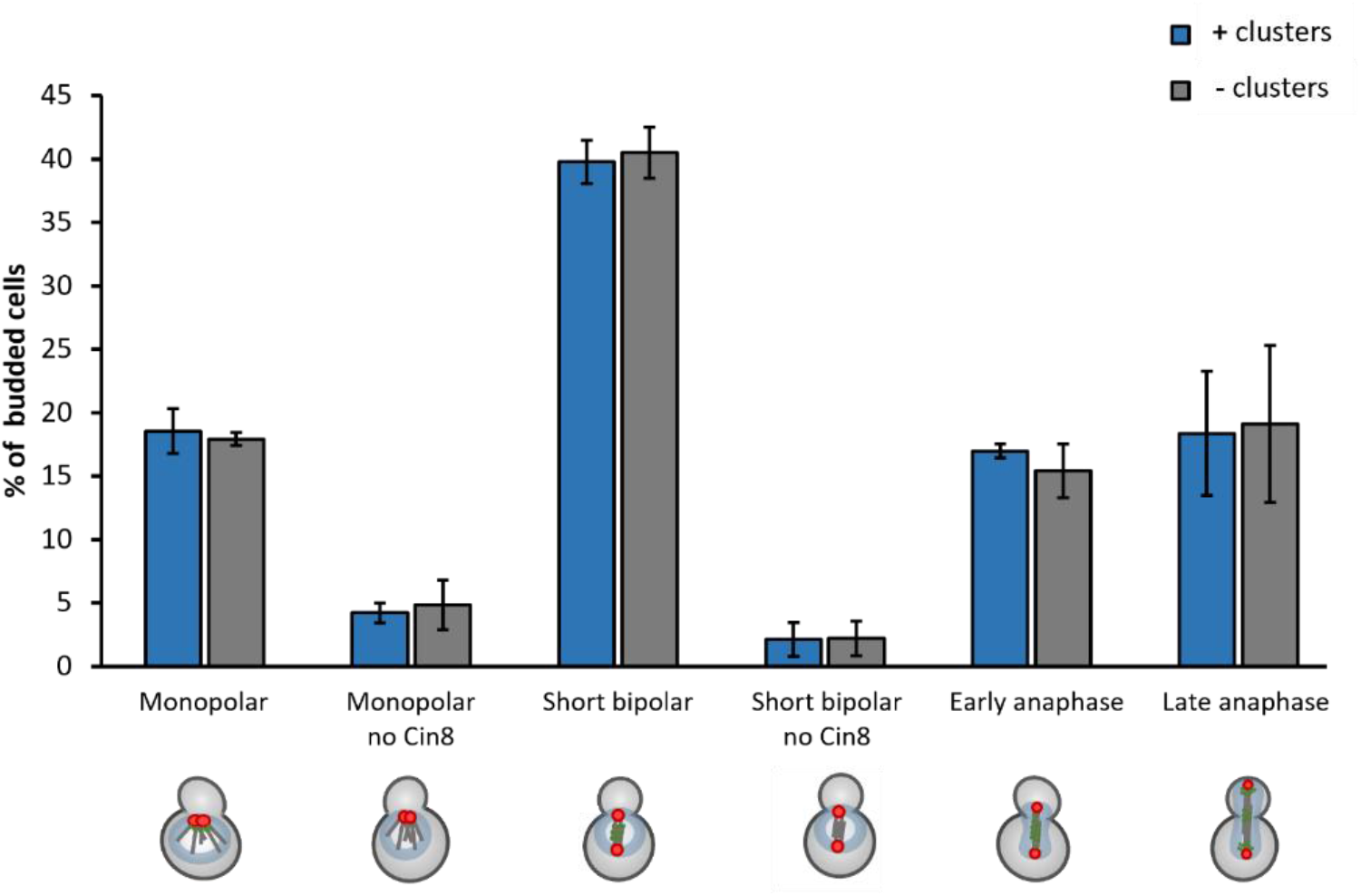
Distribution of Cin8 localization phenotypes. Comparison between two analyses: including (blue) and excluding (gray) cells in clusters. Columns and bars represent averages (± SEM) of 3 independent experiments.

These results highlight the enhanced throughput and robustness of the automated approach. In addition, we partially attribute the discrepancy between SPC tool-based and manually assessed phenotypes to inaccuracies in the manual determination of SPB positions, which directly affect the measured spindle length. In the SPC pipeline, SPB positions are automatically derived from DoG-detected maxima and then validated using a disk-ring Z-score measure, providing a precise and reproducible determination. By contrast, manual SPB selection is subjective and prone to systematic under- or overestimation, depending on the person performing the measurement. We therefore conclude that the automated, SPC-derived SPB positions (and corresponding spindle lengths) are more accurate than manual measurements and provide a more reliable basis for comparing spindle phenotypes between wild-type and mutant cells.

Overall, our results demonstrate that the SPC tool achieves high accuracy in phenotype determination while maintaining efficiency and minimizing user bias.

## Discussion

The SPC tool was developed to enable high-throughput, unbiased quantification of cell cycle phases and mitotic spindle phenotypes from three-channel 3D microscopy datasets. By integrating 2D dynamic-range projections, dual-model Cellpose deep learning for cell segmentation, and a deterministic decision tree based strictly on physical metrics, the pipeline reliably automates phenotype scoring while extracting high-dimensional spatial parameters. These include automated pole-to-pole distances (spindle lengths), spindle-bound protein coordinates, and exact bud-to-mother diameter ratios. This combination of deep-learning segmentation with rule-based physical classification fundamentally minimizes investigator-dependent analytical variability.

When validated against previously published, fully manual analyses of three *S. cerevisiae* colonies expressing wild-type Cin8 (Pandey, et al., 2021), the automated pipeline achieved comparable phenotypic distributions in a fraction of the time. Manual processing that historically required weeks or months of intensive annotation was reduced to just 1 - 2 days per colony, yielding a dramatic acceleration in experimental throughput without sacrificing analytical accuracy. Crucially, the phenotypic distributions obtained with and without interactive user correction were nearly identical (Fig. 4B). This high level of consistency demonstrates that fully automated execution alone is robust enough to accurately identify biological trends, meaning user intervention is only required for a minimal subset of borderline cases.

The remaining discrepancies between the automated and user-corrected outputs stem from two well-defined, conservative detection thresholds designed to minimize false positive results: a ∼6% baseline error rate in SPB detection and an ∼11% error rate in motor structure segmentation (Fig. 4A). While motor misdetections typically result in minor misclassifications between telophase and early/late anaphase that are rapidly corrected via the interactive interface, SPB misdetections require manual track verification.

Most importantly, SPC-derived SPB positions and spindle length determinations are less biased than traditional manual measurements. During early spindle assembly, when duplicated SPBs are separated by sub-micron distances near or below the optical diffraction limit, overlapping fluorescence emission profiles make manual coordinate selection highly subjective. Human operators may overestimate these close-proximity distances, artificially increasing anaphase scores and masking subtle metaphase delays or mid-anaphase pauses characteristic of mitotic regulator protein mutants. The SPC tool solves this limitation by using a Difference-of-Gaussians fit verified by a disk-ring Z-score, providing a precise, reproducible coordinate determination that eliminates human overestimation bias.

An additional major advantage of the SPC tool over manual annotation is its unique dual-segmentation strategy, which robustly isolates individual cell boundaries within densely packed microcolonies and cell clusters. Human annotators routinely exclude tightly clustered cells from analysis due to the visual complexity of manual tracing, introducing a systematic sampling bias toward isolated, unrepresentative cells. In our validation datasets, approximately 50% of all analysed cells belonged to dense clusters. By explicitly comparing the phenotypic distributions obtained when including or excluding clustered cell populations, we confirmed that analysing cells within dense microcolonies yields identical biological accuracy while dramatically increasing data recovery (Fig. 6).

Taken together, these results demonstrate that the SPC tool does not merely accelerate data processing; it actively enhances the accuracy, reproducibility, and mathematical reliability of quantitative spindle profiling. By removing observer bias and recovering massive pools of cluster data, this open-source pipeline provides a robust and standardized platform for executing large-scale phenotypic screens of the *S. cerevisiae* mitotic machinery.

## Acknowlegennts

This work was supported by the Israel Science Foundation (ISF) grant #629-23 awarded to LG

1 https://github.com/BIOP/ijl-utilities-wrappers; https://github.com/BIOP/ijp-LaRoMe

